# Hypersensitivity of MAIT cells to Leukocidin ED indicates a *Staphylococcus aureus* immune evasion mechanism targeting the innate effector cell response

**DOI:** 10.1101/2021.07.15.452453

**Authors:** Caroline Boulouis, Edwin Leeansyah, Srikanth Mairpady Shambat, Anna Norrby-Teglund, Johan K. Sandberg

**Author notes:** Shared last authorship (Anna Norrby-Teglund, Johan K. Sandberg). Correspondence: Dr. Johan K. Sandberg, Center for Infectious Medicine, Department of Medicine, Karolinska Institutet, 14152 Stockholm, Sweden.

## Abstract

Mucosa-associated invariant T (MAIT) cells recognize bacterial riboflavin metabolite antigens presented by MR1 and play an important role in antimicrobial immune defense. *Staphylococcus aureus* is a pathobiont expressing a range of virulence factors including the secreted toxin Leukocidin ED (LukED), which binds to certain chemokine receptors and causes cell death by osmolysis. Here, we investigated the effect of LukED on subsets of human T cells and NK cells that are involved in the early innate response to infection. MAIT cells were strikingly hypersensitive to LukED-mediated lysis and rapidly lost from the peripheral blood T cell pool upon exposure to the toxin, leaving a T cell population devoid of MAIT cells. The cytolytic effect of LukED on MAIT cells was rapid, occurred at lower LukED concentration compared to effects on the overall T cell pool, and coincided with extraordinarily high and uniform expression of CCR5. Furthermore, loss of MAIT cells was efficiently inhibited by the CCR5 inhibitor Maraviroc. Interestingly, pre-activation of MAIT cells with IL-12 and IL-18 also partially rescued these cells from LukED toxicity. Among NK cells, LukED targeted the more mature and cytotoxic CD57+ NK cell subset in a CXCR1-dependent manner. Overall, these results indicate that LukED efficiently eliminates cells of the human immune system that have the capacity to respond rapidly to *S. aureus* in an innate fashion, and that MAIT cells are exceptionally vulnerable to this toxin. Thus, the findings support a model where LukED functions as a *S. aureus* immune evasion mechanism to avoid recognition by the rapid cell-mediated responses mediated by MAIT cells and NK cells.

## Introduction

Mucosa-Associated Invariant T (MAIT) cells belong to the broad and diverse group of unconventional non-MHC restricted T cells [1, 2]. MAIT cells recognize non-peptide microbial antigens presented in complex with the MHC-class Ib-related protein (MR1) [3, 4]. Antigens recognized by MAIT cells include intermediates of the vitamin B2 (riboflavin) synthesis pathway expressed by many microbes, and this allows immune surveillance of a broad range of microorganisms in an MR1-restricted fashion [5, 6]. The presence of MAIT cells in high numbers across blood and mucosal tissues poise them to rapid effector responses in response to microbial infection [7]. Upon recognition of antigen presented by MR1 they secrete cytokines including IFNγ, TNF, IL-17A, and IL-22 [7, 8], and can participate in tissue repair and wound healing [9–12]. Furthermore, MAIT cells can kill bacterially infected cells via the release of cytotoxic effector molecules such as Granzyme A, Granzyme B, and Granulysin [13–16], and have direct antimicrobial properties against both cell-associated and free-living bacteria [17]. Their role in defense against bacterial infection was demonstrated in several mouse models of infection [18–20]. In humans, MAIT cells expand in response to *Salmonella enterica* subsp. *enterica* serovar Paratyphi A challenge [21], and migrate to lung tissue during tuberculosis [22, 23].

NK cells are the prototypical innate effector cells and mediate rapid immune responses against microbes and tumor cells [24, 25]. Differentiation of CD56^bright^ NK cells to mature cytolytic NK cells is characterized by lowered CD56 expression and the gain of CD16, CD57 and KIR expression [26, 27]. During bacterial infection, NK cells can be indirectly activated by cytokines or through interaction with other cell types, and can also directly recognize bacteria through Toll-like receptor (TLR) sensing [28]. Recently, the activating receptor KIR2DS4 was found to recognize a bacterial HLA-C*05:01-presented epitope derived from a protein conserved in many bacterial species including *S. aureus* [29].

*S. aureus* is a bacterial pathobiont that colonizes 30% of the human population through nasal and skin carriage [30]. The shift from commensal microbe to pathogen requires the expression of virulence factors as well as barrier breach [31, 32]. *S. aureus* affects local tissue and can spread systemically to cause life-threatening diseases such as pneumonia, endocarditis, and sepsis. Virulence factors that are critical for pathogenesis include superantigens, cytolytic peptides and pore-forming toxins [33]. One of the pore-forming toxins is the leucocidin ED (LukED), which is expressed by a majority of *S. aureus* isolates [34]. This bicomponent toxin is composed of two water-soluble monomers and acts in two steps: LukE first binds to target proteins on the cell membrane and recruits the LukD subunit. The complex then oligomerizes, inserts in the cell membrane as a β-barrel pore, leading to disruption of the cellular osmotic balance and cell death [34]. LukED binds to the chemokine receptors CCR5, expressed on macrophages, dendritic cells (DC) and T cells [35]. It also binds to CXCR1 and CXCR2 expressed on NK cells and neutrophils [36], as well as to DARC expressed on erythrocytes and endothelial cells [37, 38]. LukED contributes to *S. aureus* pathogenesis *in vivo* [35–37, 39], and the binding to DARC expressed on epithelial and endothelial cells leads to vascular leakage and organ failure [38]. *S. aureus* ΔLukED mutants are less invasive with reduced bacterial burden and mortality [36, 37, 39].

The role of MAIT cells in *S. aureus* immunopathogenesis remains unclear. MAIT cells are activated by *S. aureus* stimulation *in vitro* and produce IFNγ [22, 23], and their frequencies are increased in tonsils and blood of individuals with *S. aureus* tonsillitis [40]. MAIT cell are also significant contributors to the massive cytokine release, “cytokine-storm”, in response to Staphylococcal enterotoxin B (SEB), but this response renders them anergic and unable to respond to further MR1-dependent stimulation [41, 42]. In the present study, we investigate the effect of LukED on MAIT cells in comparison with other subsets in the peripheral blood T cell pool and dissect the effect of LukED on MAIT cell recognition of *S. aureus*. Furthermore, we investigate how LukED affects the NK cell compartment. Altogether, the findings indicate that LukED secretion by *S. aureus* constitutes an immune evasion mechanism to interfere with responses mediated by human innate effector cells.

## Results

### MAIT cells are hypersensitive to LukED cytotoxicity

To investigate the effect of the LukED toxin on human peripheral blood T cells, we incubated peripheral blood mononuclear cells (PBMC) in the presence of the recombinant toxins LukE and LukD and assessed population changes using flow cytometry. The total lymphocyte population was first analyzed using Uniform Manifold Approximation and Projection (UMAP) analysis [43] of healthy donor PBMC exposed to LukED. Cell populations were identified by the projection of defining markers on the UMAP topography (Figure 1A). Projection of LukED-treated versus untreated conditions revealed differences between the two settings and the loss of some cell populations (Figure 1B). Strikingly, the UMAP area defined by the 5-OP-RU-hMR1 tetramer was almost completely absent after exposure to LukED, suggesting that exposure to the toxin depletes MAIT cells. The profound reduction of MAIT cells (defined as CD3+, CD161^high^, 5-OP-RU-hMR1+) was confirmed both as percentage and absolute count (Figure 1C). In contrast, non-MAIT T cell populations were only slightly affected by the toxin with a decline of 25% (Figure 1D), versus 97% for MAIT cells (Figure 1E). The single components of the toxin alone, LukE and LukD, did not affect the T cell compartment in a detectable manner (Suppl. Figure 1A and 1B). Since LukED was previously shown to lyse T cells in a CCR5-dependent fashion (30), we zoomed in on the CCR5+ non-MAIT T cell subset. We noticed a decrease of this population (Figure 1F), which was 80% depleted upon LukED exposure (Figure 1G) compared to 97% for MAIT cells. This is in line with a strong CCR5-dependency, as MAIT cells have homogenous expression of CCR5 (Suppl. Figure 1C) and display higher level of expression compared to CCR5+ non-MAIT T cells (Figure 1H and Suppl. Figure 1D).

**Figure 1.**
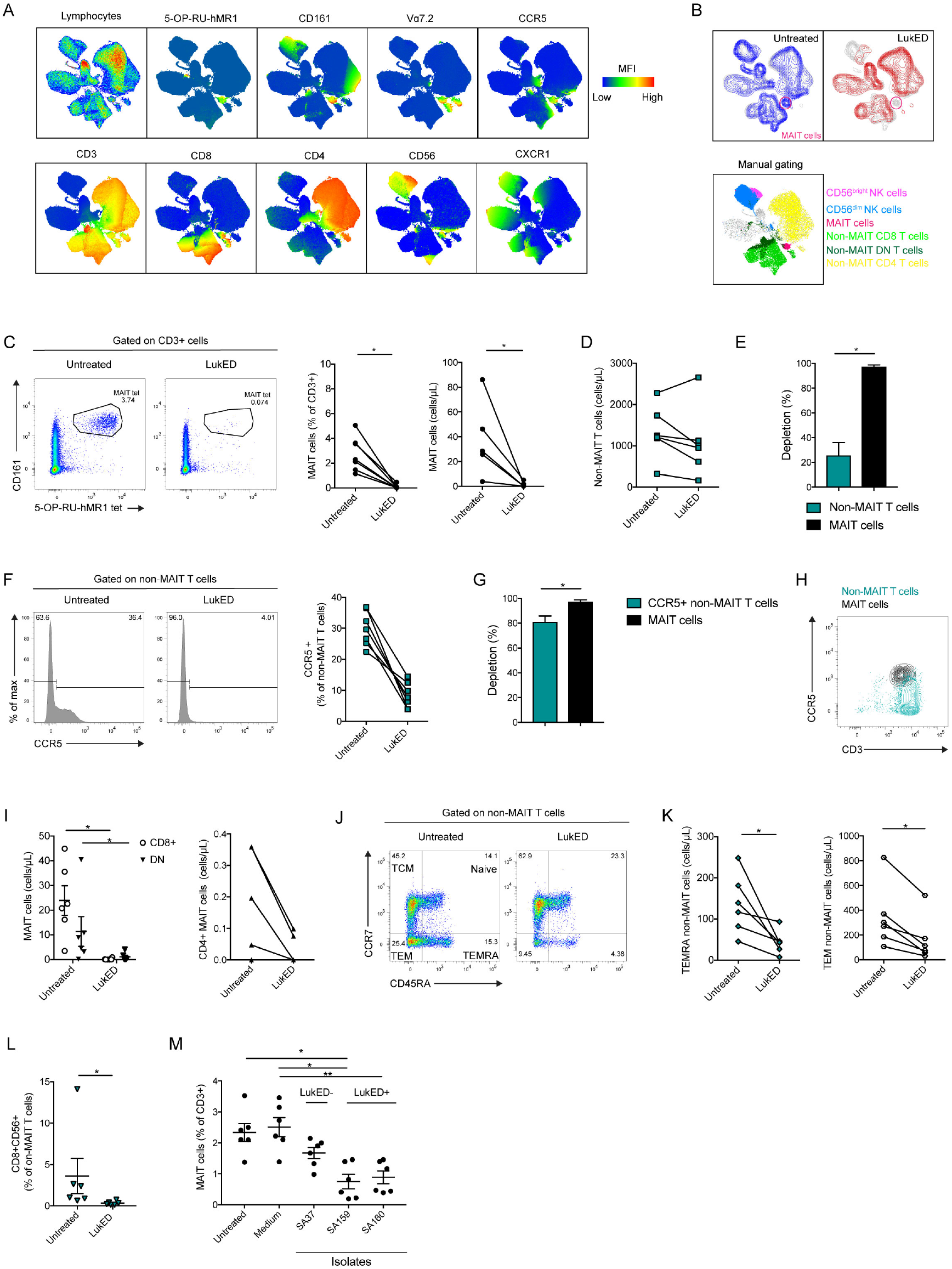
MAIT cells are the main LukED targets within the T cell pool. (A) UMAP plots of total live blood lymphocytes showing expression of the indicated markers. (B) UMAP plots of total lymphocytes exposed to LukED or not (n=6). Bottom: Overlay of immune subset identified by manual gating on the UMAP of total lymphocytes. (C) Representative flow cytometry plots, mean percentage and absolute counts of MAIT cells upon LukED treatment or not (n=6-7) (D) Absolute counts of non-MAIT T cells upon LukED exposure or not (n=6). (E) Percentage depletion upon LukED exposure between MAIT cells and non-MAIT T cells (n=6). (F) Representative flow cytometry histograms, and mean percentages of expression of CCR5+ non-MAIT T cells upon LukED treatment or not (n=6). (G) Percentages of depletion upon LukED exposure between MAIT cells and CCR5+ non-MAIT T cells (n=6). (H) Representative flow cytometry plot of CCR5 expression in MAIT cells and non-MAIT T cells (n=6). (I) Absolute counts of CD8+, CD4+ and DN MAIT cells upon LukED treatment or not (n=6). Representative flow cytometry plot of CCR7 and CD45RA expression (J) and absolute counts (K) of non-MAIT T cells with or without LukED (n=6). (L) Percentages of CD8+CD56+ non-MAIT T cells with or without LukED (n=6). (M) MAIT cell percentage upon exposure to clinical strains supernatants or medium alone (n=6). LukED was used at 5 μg/mL in the entire figure 1 and incubated for 4 h. *P<0.05, **P<0.005. The Wilcoxon’s signed-rank test was performed to detect significance in (C), (E), (G), (I), (K), (L). The Friedman test followed by Dunn’s post hoc test was used to determine significant differences between multiple, paired samples in (M). The lines and error bars represent mean and standard error.

Within the MAIT cell compartment, CD8+ and CD8-CD4-double negative (DN) MAIT cells appeared to be slightly more sensitive to LukED than the minor CD4+ MAIT cell subpopulation (Figure 1I), probably due to their relatively higher expression of CCR5 (Suppl. Figure 1E). No significant difference was noted in CD8+, CD4+ and DN non-MAIT T cells upon LukED exposure (Suppl. Figure 1F). Next, we explored the composition of the non-MAIT T cell population based on the expression of differentiation markers CCR7 and CD45RA. Terminally differentiated effector memory RA+ (TEMRA) and effector memory T (TEM) cell populations decreased in frequency upon LukED exposure (Figure 1J and 1K) contrary to central memory (TCM) and naïve T cells (Suppl. Figure 1G). This coincided with higher levels of CXCR1 and CCR5 expression by TEMRA cells and TEM cells, respectively (Suppl. Figure 1H). Projection of the LukED-treated condition on the UMAP also revealed a dramatic loss of cells double expressing CD8 and CD56 (Figure 1B). The decrease of CD8+CD56+ non-MAIT T cells was confirmed by manual gating (Figure 1L), and coincided with co-expression of CCR5 and CXCR1 on this subset (Suppl. Figure 1K). Overall, the loss of TEMRA, TEM and CD8+CD56+ non-MAIT T cells was not as severe as the depletion of MAIT cells (Suppl. Figure 1I and 1L).

To investigate if the patterns obtained with the recombinant toxin occur with live bacteria, we incubated PBMC with culture supernatants of three *S. aureus* clinical isolates previously characterized regarding their toxin gene profile [44]. The three strains studied were selected based on varying presence of the *luk*ED genes and low expression of the bacterial α-toxin, another pore-forming toxin highly effective in killing cells by binding the surface protein ADAM-10. Interestingly, PBMC cultures exposed to the supernatants of *luk*ED positive strains lost a larger fraction of their MAIT cells, as compared to cultures exposed to *luk*ED negative strains (Figure 1M). CCR5+ non-MAIT T cells followed a similar pattern (Suppl. Figure 1M). Taken together, these findings indicate that LukED depletes T cell subsets with effector and effector memory characteristics, with MAIT cells being the major targeted population.

### Maraviroc rescues MAIT cells from LukED toxicity

To investigate if LukED toxicity against MAIT cells is CCR5-dependent, we evaluated the ability of Maraviroc (MVC), a CCR5 antagonist used in HIV therapy, to protect MAIT cells from the recombinant toxin. Addition of MVC to the assay largely rescued the MAIT cell population from LukED toxicity (Figure 2A). While this protective effect was incomplete, it was observed for all the main CD8, CD4 and DN MAIT cell subsets (Suppl. Figure 2A). MVC did not have a detectable effect on the overall non-MAIT T cells bulk population (Figure 2B), but seemed to partially rescue the CCR5+ subset of non-MAIT T cells (Figure 2C). Similarly, there was a trend towards partial MVC rescue of TEMRA, TEM and CD8+CD56+ non-MAIT T cells but this effect did not reach statistical significance (Suppl. Figure 2B and 2C). As some of those subsets express CXCR1, LukED binding to this receptor would not be expected to be inhibited by MVC. Altogether, these results indicate that MVC inhibits the toxicity of LukED against MAIT cells *in vitro*.

**Figure 2.**
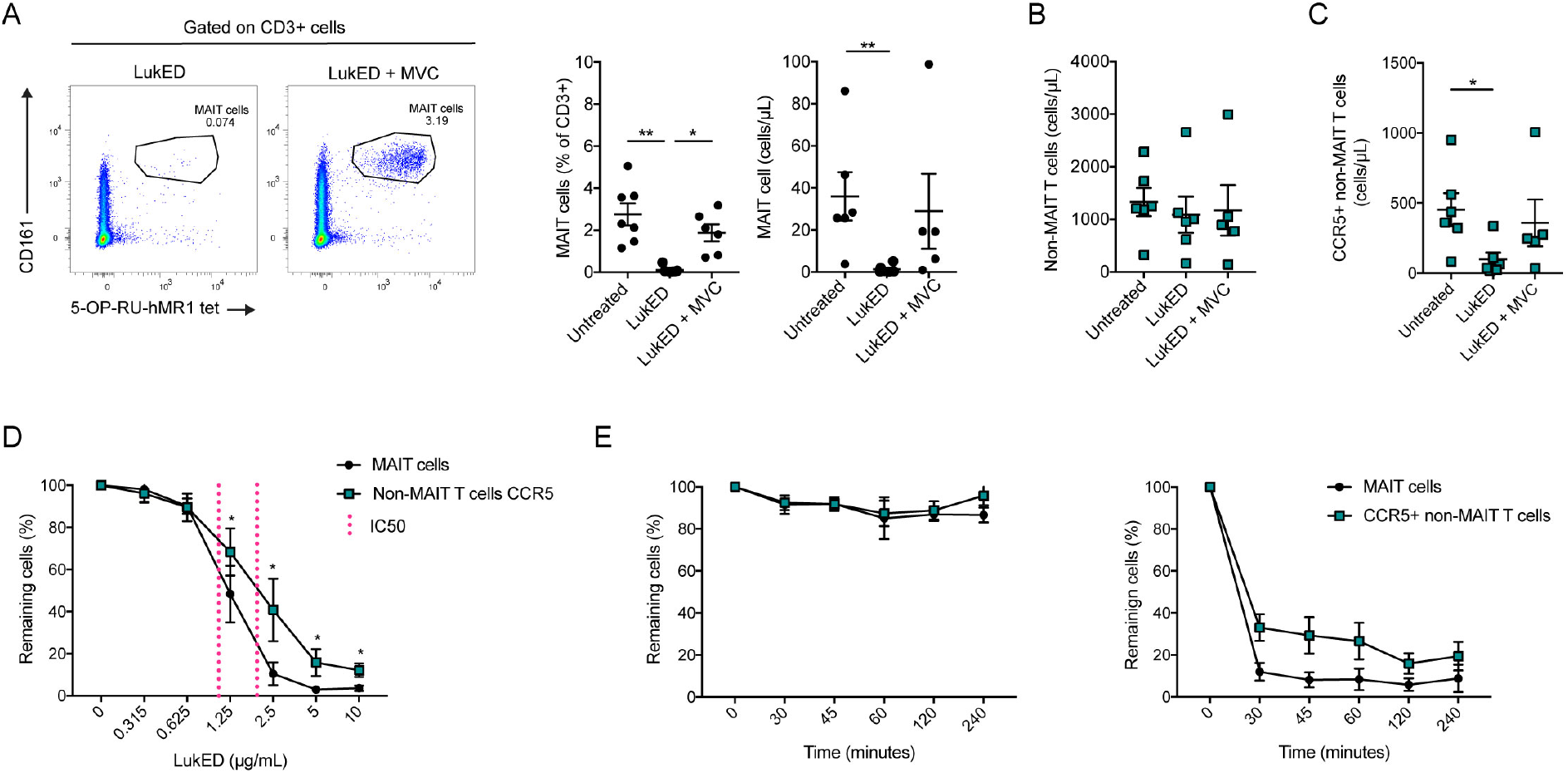
Maraviroc rescues MAIT cells from LukED *in vitro*. (A) Representative flow cytometry plots, mean percentage, and absolute count of MAIT cells upon LukED treatment with or without MVC (n=6-7). MVC is used at 1 μg/mL. LukED was used at 5 μg/mL for 4 h in (A), (B) and (C). (B) Absolute cell counts of non-MAIT T cells upon LukED exposure with or without MVC (n=5-6). (C) Absolute counts of CCR5+ non-MAIT T cells upon LukED exposure with or without MVC (n=5-6). (D) Relative percentages of MAIT cells and CCR5+ non-MAIT T cells over a range of LukED concentrations after 2 h of exposure. The 50% maximal inhibitory concentration (IC50) is indicated in magenta (n=6). (E) Relative percentages of MAIT cells and CCR5+ non-MAIT T cells after exposure of low dose of LukED (0.625 μg/mL; left) or high dose of LukED (5 μg/mL; right) from 30 min to 4 h (n=6). *P<0.05, **P<0.005. The Kruskal-Wallis test followed by Dunn’s post hoc test was used to detect significant differences in (A) and (C). The Wilcoxon’s signed-rank test was performed to detect significance between the two populations in (D). The lines and error bars represent mean and standard error.

### LukED depletes MAIT cells with rapid kinetics and at low doses

We next evaluated the dose sensitivity and kinetics of action of LukED on MAIT cells and CCR5+ non-MAIT T cells. Titration curves revealed dose-dependent depletion of MAIT cells, as well as of CCR5+ non-MAIT T cells (Figure 2D). Interestingly, at the dose range from 1.25 μg/mL to 10 μg/mL, MAIT cells were more sensitive than CCR5+ non-MAIT T cells. Furthermore, the amount of LukED needed to kill half of the population (IC50) was lower for MAIT cells (1.09 μg/mL) than for CCR5+ non-MAIT T cells (2.17 μg/mL) (Figure 2A, magenta reference line). Thus, MAIT cells are significantly more vulnerable to LukED at intermediate doses. Depletion occurred rapidly, within 30 min of incubation, and longer duration of LukED exposure did not substantially increase MAIT cell and CCR5+ non-MAIT T cell depletion (Figure 2E). Altogether these findings showed that the lethal effect of LukED on MAIT cells is rapid, dose-dependent, and occurs at lower doses compared to conventional T cells.

### MAIT cell activation with IL-12 and IL-18 partly prevents LukED-mediated loss

MAIT cells can be activated in response to innate cytokines produced in the setting of myeloid cell activation, such as IL-12 and IL-18. Surprisingly, MAIT cells activated by IL-12 and IL-18 for 20 h, and then exposed to LukED for the last two hours of incubation, were partly preserved as compared to the non-activated control MAIT cells (Figure 3A and 3B). This effect coincided with a decrease in CCR5 expression upon IL-12 and IL-18 stimulation (Figure 3C). It is thus possible that activation by innate cytokines in the inflammatory milieu may render MAIT cells partly resistant to LukED.

**Figure 3.**
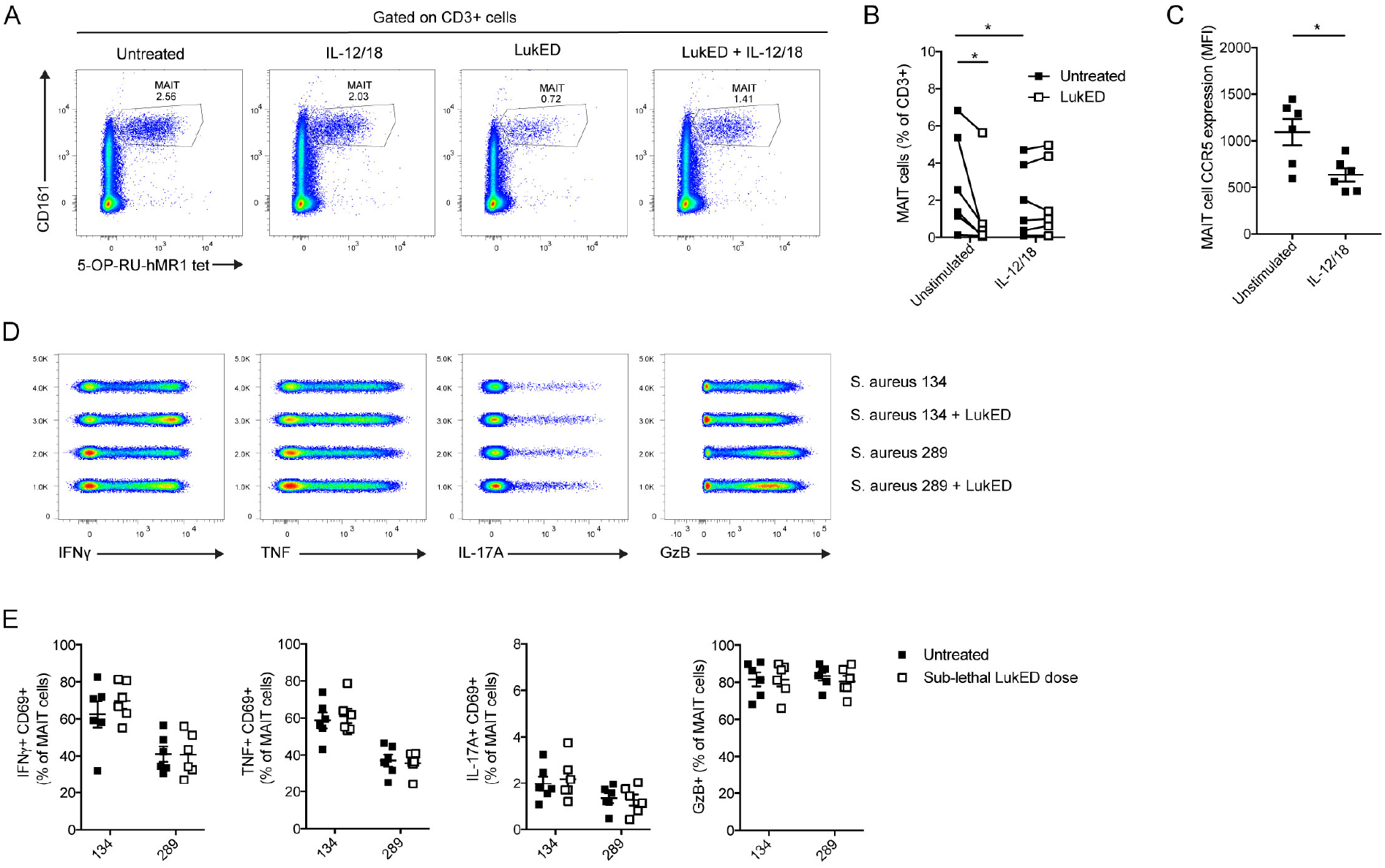
MAIT cell activation with innate cytokines makes them partly resistant to LukED. Representative flow cytometry plots (A) and levels (B) of MAIT cells following 20 h stimulation with IL-12 and IL-18, in the presence or absence of LukED at 1.25 μg/mL during the last 2 h of stimulation (n=6). (C) CCR5 expression on MAIT cells stimulated 18 h with IL-12 and IL-18 (n=6). Concatenated flow cytometry plots (D) and combined data (E) of the expression of IFNγ+CD69+, TNF+CD69+, IL-17A+CD69+ and GzB+ MAIT cells untreated or stimulated with THP-1 fed *S. aureus* 134 (LukED negative) or 289 (LukED positive) for 24 h at the microbial dose of 30, after exposure to a sublethal dose of LukED (0.312 μg/mL). *P<0.05. The Wilcoxon’s signed-rank test was performed to detect significance in (B) and (C). The lines and error bars represent mean and standard error.

### Sub-lethal doses of LukED have no major impact on MAIT cell responsiveness

Since LukED is a pore-forming toxin it is possible that it may affect signaling in MAIT cells exposed to concentrations insufficient for direct killing of the cells. We therefore next sought to determine if exposure to sub-lethal doses of LukED affects TCR-dependent activation of MAIT cells. To test this, MAIT cells were pre-treated with sub-lethal doses of LukED before stimulation with the mildly fixed LukED-positive *S. aureus* strain 134, or the LukED-negative strain 289. No detectable difference was observed in MAIT cell IFNγ, TNF, IL-17A and GzB production between the sub-lethal dose LukED-treated and -untreated conditions (Figure 3D and 3E). These results indicate that sub-lethal levels of LukED have little observable impact on MAIT cell function and ability to respond to bacteria.

### LukED targets the mature CD57+ NK cell subset

In the UMAP topography (Figure 1A and 1B), apart from MAIT cells another area severely affected by LukED was the one dominated by CD56+CD3- cells corresponding to the NK cell population. Indeed, LukED severely reduced both CD56^bright^ and CD56^dim^ NK cells among PBMC both as percentage and as absolute count (Figure 4A), and the CD56^dim^ subset was particularly severely impaired (Figure 4B). LukE and LukD alone did not affect NK cells (Suppl. Figure 3A). In line with the differential activity of LukED on the two NK cell subsets, CD56^dim^ NK cells had higher expression of CXCR1 and CXCR2 than CD56^bright^ NK cells (Figure 4C and 4D). In general, NK cells expressed low levels of CCR5 (Figure 4D and Suppl. Figure 3B). Dissection of the NK cell compartment revealed that LukED targeted the most mature NK cells known to have cytolytic effector properties. In particular, the toxin diminished CD56^dim^ NK cells expressing CD16, CD57, KIRD2L1, perforin or co-expressing CD57 and KIRD2L1 (Figure 4E and 4F). LukED exposure did not change the representation of NK cell populations expressing granulysin or CD27 (Suppl. Figure 3C). Taken together, these observations indicate that LukED preferentially targets mature cytolytic NK cells.

**Figure 4.**
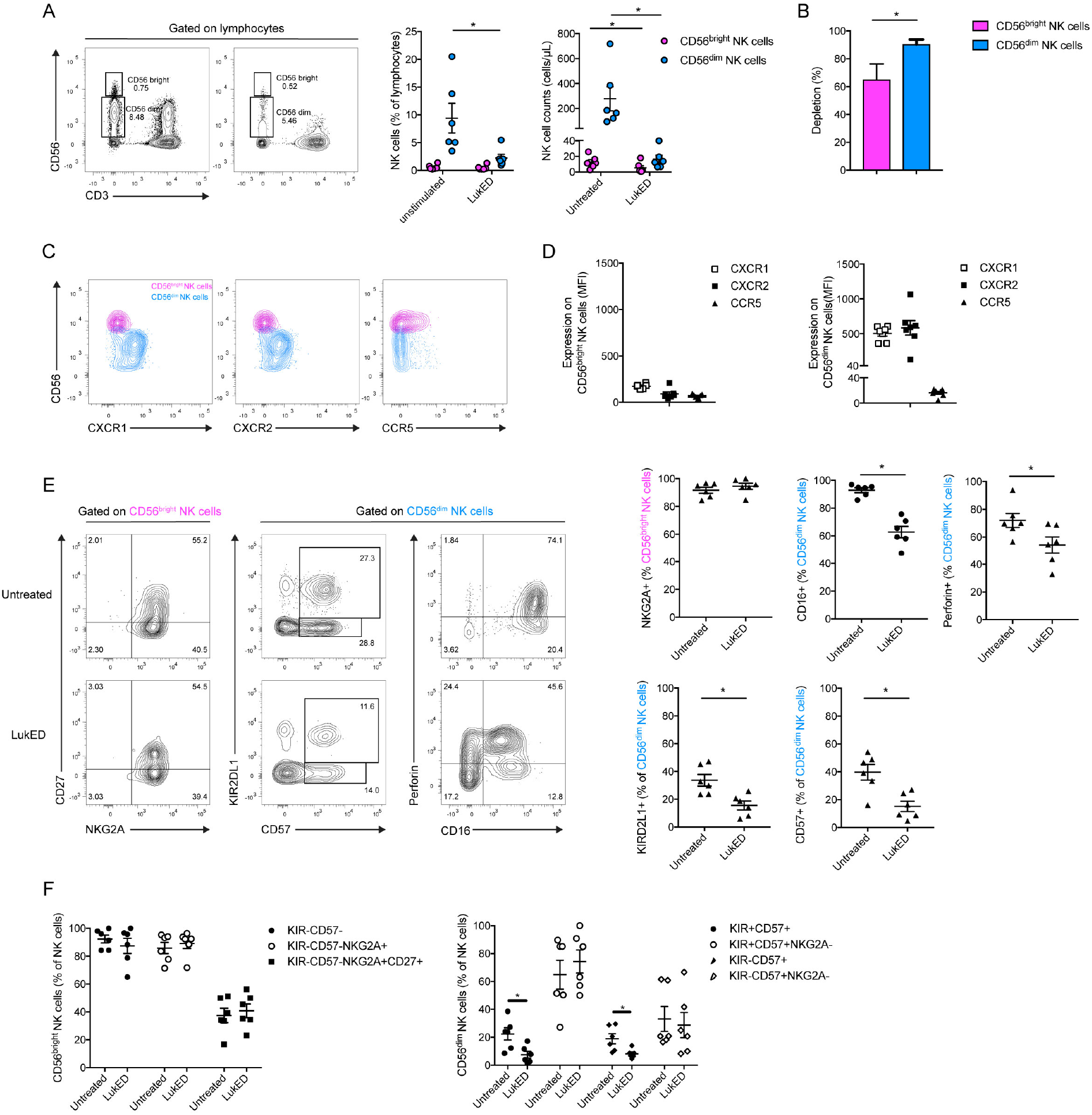
LukED targets mature CD56^dim^ NK cells. (A) Representative flow cytometry plots and combined donor data, of the percentage and absolute count of CD56^bright^ and CD56^dim^ NK cells upon LukED treatment for 4 h (n=6). LukED is used at 5 μg/mL throughout the figure. (B) Percentage of depletion of CD56^bright^ and CD56^dim^ NK cells upon LukED treatment (n=6). Representative flow cytometry plots (C) and combined data (D) of the expression of CXCR1, CXCR2 and CCR5 on CD56^bright^ and CD56^dim^ NK cells (n=6). (E) Representative flow cytometry plots and combined data of the expression of CD27 and NKG2A on CD56^bright^ and the expression of KIRD2L1, CD57, Perforin and CD16 on CD56^dim^ NK cells with or without LukED treatment (n=6). (F) Combined data of the percentage of co-expression of the indicated markers on CD56^bright^ (left) and CD56^dim^ (right) NK cells with or without LukED exposure (n=6). *P<0.05. The Wilcoxon’s signed-rank test was performed to determine significance. The lines and error bars represent mean and standard error.

## Discussion

The soluble virulence factors secreted by *S. aureus* play a major role in the pathogenesis of this infection. However, it is still not fully understood how these virulence factors affect the human immune system. MAIT cells are an attractive target for development of vaccines and therapeutics due to their specificity towards conserved bacterial antigen, their secretion of IFNγ, IL-17, and IL-22, as well as their presence in skin and blood, the main sites of *S. aureus* infection and pathogenesis [45]. In this study, we explored the interplay between the LukED toxin and the peripheral blood immune system, and found that MAIT cells are hypersensitive to LukED, due to their very high surface expression of CCR5. LukED depleted MAIT cells more efficiently, with lower IC50, than other T cells subsets, such as conventional TEM and TEMRA cells. The mature effector CD56^dim^ NK cells were also depleted to a large extent, while CD56^bright^ NK cells were less affected.

MAIT cells and NK cells are important effector cells involved in the host response against bacterial infection. MAIT cells have cytotoxic properties and can directly kill bacteria [14, 15, 17]. NK cells are activated by *S. aureus* [46] and contribute to immune defense against this infection *in vivo* [47]. In the absence of Th1 and Th17 immunity the susceptibility to *S. aureus* infection is increased [45]. Thus, the findings of the present study indicate that LukED targets cells that have the capacity to mount a rapid effector response against *S. aureus*. In particular, the rapid and almost complete elimination of MAIT cells by LukED may represent an important immune evasion mechanism affecting the outcome of *S. aureus* infection. Interestingly, recent findings indicate that pathogenic *Salmonella* Typhimurium sequence type 313 strains escape MAIT cell recognition through overexpression of the RibB enzyme, thereby severely limiting the production of MR1-presented antigen [48]. Thus, emerging evidence indicates that several distinct species of bacterial pathogens have evolved mechanisms to avoid innate immune control by MAIT cells.

The CCR5 antagonist MVC inhibits the effect of LukED on T cells. As MVC is approved for use in humans, it is possible that this inhibitor could be used to block MAIT cell depletion and restore immune control in patients with invasive *S. aureus* infection. It is interesting to note that several new treatments are in development against *S. aureus* infection including monoclonal antibodies aimed to neutralize *S. aureus* toxins [49]. In experimental animal models of infection in mice [50] and rabbits [51], the combination of monoclonal antibodies, including one targeting LukED, shows enhanced efficacy over single antibody administration with decreased bacterial burden [50] or increased survival [51]. In addition, new neutralizing agents called centyrins are able to block the binding of the five bi-component leukocidins to their receptors and protect *in vivo* against *S. aureus* infection [52].

*S. aureus* is a prominent pathogen in the hospital setting and the treatment is complicated by the emergence of methicillin-resistant *S. aureus* (MRSA) with acquisition of a multiresistance profile [32, 53]. MAIT cells were recently shown to recognize antibiotic resistant bacteria and their effector functions can overcome certain bacterial resistance mechanisms [17]. Furthermore, our findings indicate that MAIT cell functionality was retained upon exposure to sub-lethal doses of LukED, supporting the notion that low concentrations of toxin in the microenvironment do not disrupt signaling in MAIT cells. In this context, neutralization of these *S. aureus* virulence factors may rescue the effector function of MAIT cells, even if neutralization is not complete.

Finally, we speculate that the downregulation of CCR5 on MAIT cells upon activation with IL-12 and IL-18 may allow MAIT cells to evade LukED toxicity. MAIT cells recruited to tissue sites in the context of innate immune activation in response to bacteria can possibly in this way be partially protected against the toxin. Perhaps this effect can be harnessed to promote a more resistant phenotype in MAIT cells. In summary, we show that the LukED toxin is a major killer of cytolytic effector immune cells in blood. Notably, MAIT cells are hypersensitive to the toxin compared to other types of cytolytic effector cells such as conventional T cells and NK cells. These findings support a model where LukED targeting of CCR5 is an *S. aureus* immune evasion mechanism to avoid rapid MAIT cell detection and elimination.

## Methods

### Ethics statement

Peripheral blood was collected from healthy adults recruited at the Blood Transfusion Clinic at the Karolinska University Hospital Huddinge. All donors gave written informed consent in accordance with study protocols conforming to the provisions of the Declaration of Helsinki and approved by the Regional Ethics Review Board in Stockholm (approval number 2016/1415-32).

### Microbes

*Staphylococcus aureus* strains 134 and 289 were cultured overnight at 37°C in CCY broth. Bacterial counts were determined by the standard plate counting method on appropriate culture media, and counts were expressed as colony-forming unit (CFU)/mL. The microbes were then stored at −80°C in 50% glycerol/50% PBS.

### Cell isolation

PBMCs were isolated from peripheral blood by Ficoll–Hypaque density gradient centrifugation (Lymphoprep, Axis-Shield). After isolation, PBMCs were rested overnight in RPMI-1640 medium supplemented with 25 mM HEPES, 2 mM L-glutamine (all from Thermo Fisher Scientific), 10% FBS (Sigma-Aldrich), 50 μg/mL gentamicin (Life Technologies), and 100 μg/mL normocin (InvivoGen) (complete medium). From PBMC, we isolated Vα7.2+ cells using anti-Vα7.2 PE-conjugated mAb (Biolegend), followed by positive selection with magnetic-activated cell sorting (MACS) and anti-PE microbeads (Miltenyi Biotec), as previously described and per the manufacturer’s instructions [16, 54].

### Supernatant assay

Clinical *S. aureus* strains were cultured overnight in CCY medium at 37°C. Supernatant of the culture was harvested by centrifugation and sterilized through 0.2 μm pore size filters to obtain the bacterial free culture supernatant. PBMC were incubated with the supernatant at 1:20 dilution for 4 h.

### LukED assays

PBMC were incubated with recombinant LukED (IBTBioservices) at the indicated concentration between 45 min and 6 h, as indicated in the text. In selected experiments, MVC (Sigma) at 1 μg/mL was added just before the recombinant LukED.

### MAIT cell activation assays

THP-1 cells were seeded in complete medium for 2 h prior pulsing with bacteria. *S. aureus* 134 and 289 were washed once in PBS before fixation in 1% formaldehyde for 3 min, and extensive PBS washes. The bacteria were then resuspended in complete medium and fed to THP-1 monocytic cells at the microbial dose of 30. Vα7.2+ cells were pre-incubated for 1 h and 30 min with LukED at 0.312 μg/mL before adding to the bacteria-pulsed THP-1 and co-culture at 2:1 ratio for 24 h in the presence of 1.25 μg/mL anti-CD28 mAb (L293, BD Biosciences). Monensin and brefeldin A (both from BD Biosciences) were added to the last 6 h of culture. The THP-1 cell line was maintained in complete medium and was routinely tested negative for mycoplasma. For TCR-independent activation, PBMC were incubated for 20 h with IL-12 (Peprotech) and IL-18 (MBL) at 10 ng/mL and 100 ng/mL, respectively. LukED at 1.25 μg/mL was added for the two last hours of culture in some conditions.

### Flow cytometry

Tetramer staining with 5-OP-RU-loaded h-MR1-PE (NIH Tetramer Core Facility) was performed at 20 min at room temperature before staining with monoclonal antibodies for other markers. Antibodies used: anti-CD3 Bv650 (OKT3, Biolegend), anti-CD3 FITC (SK7, BD Biosciences), anti-Vα7.2 PE (3C10, Biolegend), anti-Vα7.2 PE-Cy7 (3C10, Biolegend), anti-CD161 Pe-Cy5 (DX12, BD Biosciences), anti-CD4 Bv711 (OKT4, Biolegend), anti-CD8 Bv570 (RPA-T8, Biolegend), anti-CD56 BUV737 (NCAM16.2, BD Biosciences), anti-CCR7 Bv421 (150503, BD Biosciences), anti-CD45RA AF700 (HI100, BD Biosciences), anti-CCR5 BUV395 (2D7/CCR5, BD Biosciences), anti-CXCR1 AF488 (8F1/CXCR1, BD Biosciences), anti-CXCR2 Bv785 (6C6, BD Biosciences), anti-CD158b Bv510 (CH-L, BD Biosciences), anti-CD16 Bv711 (3G8, BD Biosciences), anti-CD57 PE-Cy5 (NK-1, BD Biosciences), anti-CD27 PE-Cy7 (M-T271, Biolegend), anti-NKG2A APC (REA110, Miltenyi), anti-CD14 APC-Cy7 (MѻP9, BD Biosciences), anti-CD19 APC-Cy7 (SJ27C1, BD Biosciences) anti-CD69 BUV737 (FN50, BD Biosciences), anti-GzB FITC (GB11, Biolegend), anti-IFNγ APC (25723.11, BD Biosciences), anti-TNF PE-Cy7 (Mab11, BD Biosciences), anti-IL-17A Bv421 (BL168, Biolegend), anti-Perforin Bv421 (B-D48, Biolegend), anti-Granulysin (DH2, BD Biosciences) LIVE/DEAD Fixable Near-IR dye (Invitrogen). Flow cytometry data was acquired on BD LSRFortessa or BD Symphony A5 instruments (both BD Biosciences) and analyzed using FlowJo software v.10.5.3 (TreeStar).

### Statistics

Statistical analyses were performed using Prism software v.7 (GraphPad). Data sets were first assessed for normality of the data distribution. Statistically significant differences between samples were determined as appropriate using the unpaired t-test or Mann-Whitney’s test for unpaired samples, and the paired t-test or Wilcoxon’s signed-rank test for paired samples. Correlations were assessed using the Spearman’s rank correlation. Two-sided p-values < 0.05 were considered significant.

## Author contributions

J.K.S. and A.N.T. designed the study.

C.B. performed experiments.

S.M.S and A.N.T. provide characterized clinical strains.

C.B., E.L., A.N.T and J.K.S. interpreted the data.

C.B. and J.K.S. wrote the paper.

All authors read and revised the manuscript.

## Acknowledgements

The MR1 tetramer technology was developed jointly by Dr. James McCluskey, Dr. Jamie Rossjohn, and Dr. David Fairlie, and the material was produced by the NIH Tetramer Core Facility as permitted to be distributed by the University of Melbourne. This research was supported by grants to JKS from the Swedish Research Council (2016-03052), the Swedish Cancer Society (CAN 2017/777), the Swedish Heart-Lung Foundation (20180675), and the Center for Innovative Medicine (20190732). Further support to EL was from the Swedish Research Council (2015-00174), and Marie Skłodowska Curie Actions, Cofund, and to ANT from Center for Innovative Medicine and Region Stockholm [20180058], Swedish Research Council (2018-02475, 2018-131 under the frame of ERA PerMED), Swedish Governmental Agency for Innovation Systems under the frame of NordForsk (90456). The funders had no role in study design, data collection and analysis, decision to publish, or preparation of the manuscript.

---

**Supplementary Figure 1.**
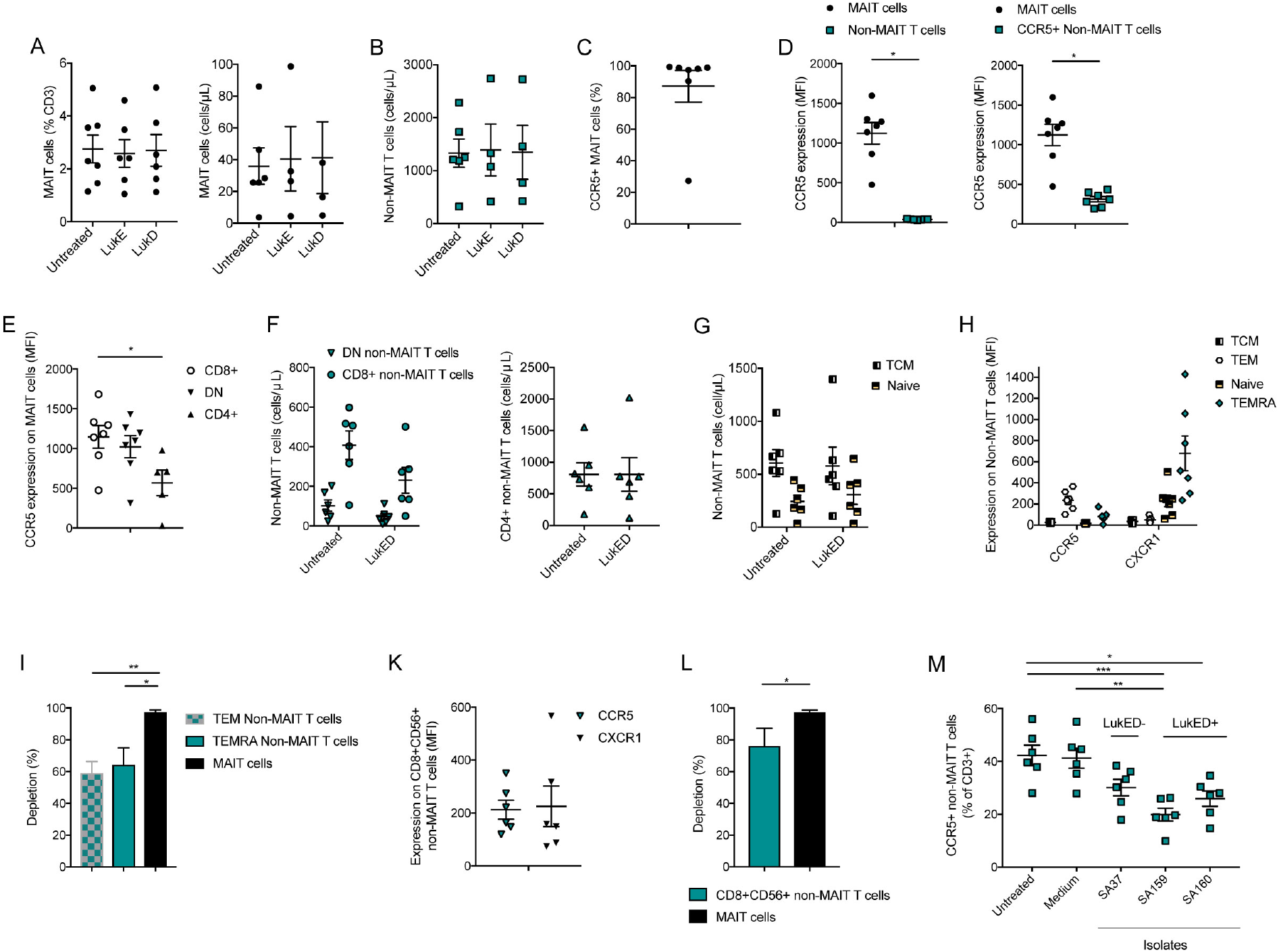
LukED effects on MAIT cells and non-MAIT T cells. (A) Combined data of the percentage (left) or absolute counts (right) of MAIT cells with or without the presence of LukE and LukD (n=4-6). (B) Absolute counts of non-MAIT T cells with or without the presence of LukE and LukD (n=4-6). (C) Percentage of CCR5+ MAIT cells (n=7). (C) CCR5 expression on MAIT cells versus non-MAIT T cells (left) or CCR5+ non-MAIT T cells (right) (n=7). (E) CCR5 expression on CD8+, CD4+ and DN MAIT cells (n=5-7). (F) Absolute counts of CD8+, CD4+ and DN non-MAIT T cells with or without LukED at 5 μg/mL for 4 h. (G) Absolute counts of TCM and Naïve non-MAIT T cells with or without LukED at 5 μg/mL for 4 h. (H) Combined data of CCR5 and CXCR1 expression on non-MAIT T cells non-MAIT T cell subsets (n=7). (I) Percentage of depletion upon LukED exposure (5 μg/mL for 4 h) of TEM, TEMRA non-MAIT cells and MAIT cells (n=6). (K) Combined data of CCR5 and CXCR1 expression on CD8+CD56+ non-MAIT T cells (n=6). (L) Percentage of depletion upon LukED exposure (5 μg/mL for 4 h) of CD8+CD56+ non-MAIT T cells versus MAIT cells (n=6). (M) CCR5+ non-MAIT T cell percentage upon exposure to clinical strains supernatants or medium (n=6). *P<0.05, **P<0.005, ***P<0.0005. The Wilcoxon’s signed-rank test was used to detect significant differences in (D). The Kruskal-Wallis test followed by Dunn’s post hoc test was used to detect significant differences in (E) and (I). Then Mann-Whitney test was applied in (L) to detect significance. The Friedman test was used to detect significance in (M). The lines and error bars represent mean and standard error.

**Supplementary Figure 2.**
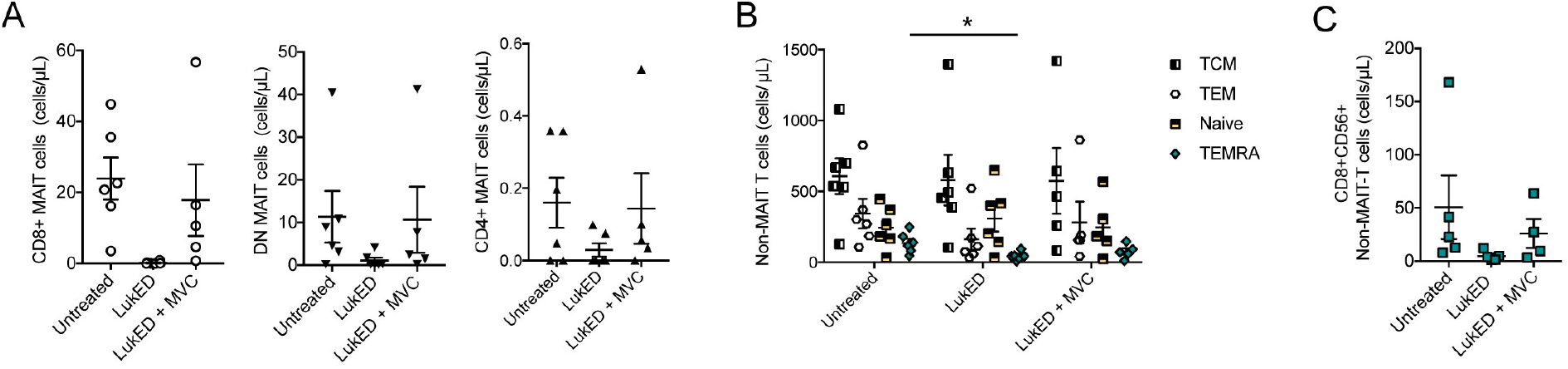
Subsets of MAIT cells are rescued by Maraviroc. (A) Combined data of the absolute count of CD8+, CD4+ and DN MAIT cells in the presence of 5 μg/mL LukED, with or without MVC (1 μg/mL) (n=5-6). (B) Absolute counts of CM, EM, TEMRA and naïve non-MAIT T cells in the presence of 5 μg/mL LukED, with or without MVC (1 μg/mL) (n=5-6). (C) Absolute counts of CD8+CD56+ non-MAIT T cells in the presence of 5 μg/mL LukED, with or without MVC (1 μg/mL) (n=5-6). *P<0.05. The Wilcoxon’s signed-rank test was used to detect significant differences in (B). The lines and error bars represent mean and standard error.

**Supplementary Figure 3.**
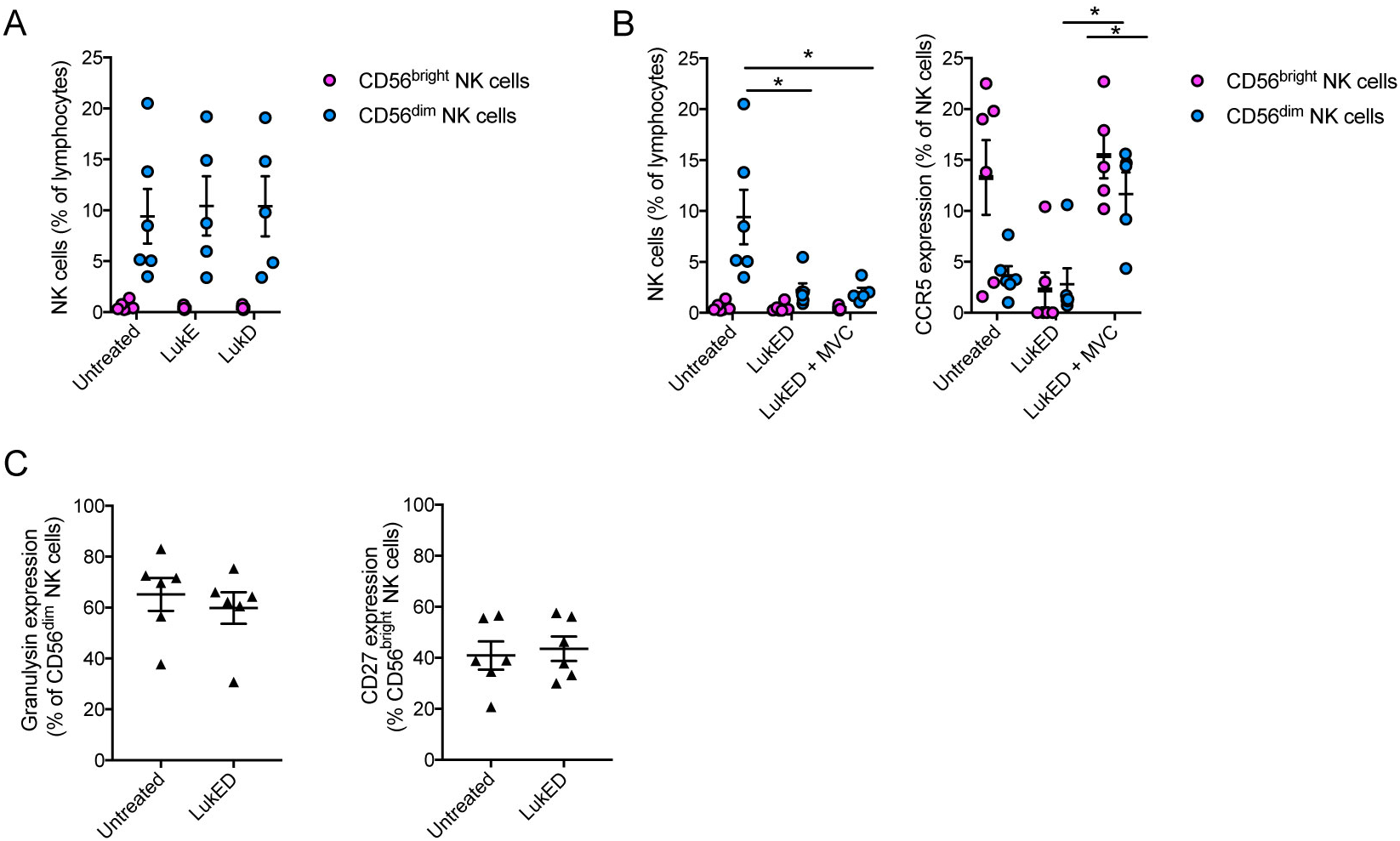
LukED targets NK cells. (A) Combined data of the percentage of CD56^bright^ and CD56^dim^ NK cells with or without the presence of LukE and LukD (n=5-6). (B) Combined data of the percentage of CD56^bright^ and CD56^dim^ NK cells (left) and of CCR5+ CD56^bright^ and CD56^dim^ NK cells (right) in the presence of LukED (5 μg/mL for 4 h), with or without MVC (1 μg/mL) (n=5-6). (C) Combined data of the expression of granulysin on CD56^dim^ NK cells and CD27 on CD56^bright^ NK cells with or without LukED (5 μg/mL for 4 h) (n=6). *P<0.05. The Kruskal-Wallis test followed by Dunn’s post hoc test was used to detect significant differences in (B). The lines and error bars represent mean and standard error.

